# Systematic optimization of siRNA productive uptake into resting and activated T cells *ex vivo*

**DOI:** 10.1101/2023.10.20.563275

**Authors:** A Kremer, T Ryaykenen, RA Haraszti

## Abstract

RNA-based medicines are ideally suited for precise modulation of T cell phenotypes in anti-cancer immunity, in autoimmune diseases and for *ex vivo* modulation of T-cell-based therapies. Therefore, understanding productive siRNA uptake to T cells is of particular importance.

Most studies used unmodified siRNAs or commercially available siRNA with undisclosed chemical modifications patterns to show functionality in T cells. Despite being an active field of research, robust siRNA delivery to T cells still represents a formidable challenge. Therefore, a systematic approach is needed to further optimize and understand productive siRNA uptake pathways to T cells.

Here we compared conjugate-mediated and nanoparticle-mediated delivery of siRNAs to T cells in the context of fully chemically modified RNA constructs. We showed that lipid-conjugate-mediated delivery outperforms lipid-nanoparticle-mediated and extracellular-vesicle-mediated delivery in activated T cells *ex vivo*. Yet, ex vivo manipulation of T cells without the need of activation is of great therapeutic interest for CAR-T, engineered TCR-T and allogeneic donor lymphocyte applications. We are first to report productive siRNA uptake into resting T cells using lipid-conjugate mediated delivery. Interestingly, we observed strong dependence of silencing activity on lipid-conjugate-identity in resting T cells but not in activated T cells. This phenomenon is consistent with our early uptake kinetics data. Lipid-conjugates also enabled delivery of siRNA to all mononuclear immune cell types, including both lymphoid and myeloid lineages. These findings are expected to be broadly applicable for *ex vivo* modulation of immune cell therapies.

## Introduction

Hyperactivation of specific immune cell subsets and/or pathways drive hematological malignancies, autoimmune diseases or underly toxicity of cell therapeutics including that of CAR-T cells. Current therapeutics often fail to specifically target disease pathways and co-inhibit numerous pathways in numerous cell subsets instead[1-3]. Approved therapies often inhibit surrogate targets when the disease specific target is undruggable[4, 5]. Both strategies lead to significant toxicities or primary treatment failure. Precision targeting, therefore, is urgently needed to enhance safety and efficacy of therapeutic applications in immune cells.

siRNAs, a novel precision medicine drug class, hold promise to meet this clinical need to selectively inhibit any sequence (including noncoding sequences[6]) in the transcriptome. However, siRNA delivery to immune cells (i.e. effector cells of most hematological and autoimmune diseases as well as in cell therapeutics) has proved challenging.

Delivery to T cells has been in the focus of several studies to date showing varying extent of siRNA uptake upon formulation into lipid nanoparticles[7], conjugation to lipids[8], aptamers[9, 10] and antibodies[11-14] or upon electroporation[15]. All of these studies used unmodified siRNAs[9-12, 15] or commercially available siRNAs with an undisclosed chemical modification pattern[7, 8, 13, 14]. One study has shown improved silencing activity of partially modified versus unmodified siRNA[16] and another study showed an advantage of fully modified compared to partially modified siRNAs in activated T cells[17]. These findings are consistent with a wealth of data now showing that full chemical stabilization of siRNAs is required for *in vivo* and clinical applications targeting liver as well as extrahepatic tissues[18]. Full chemical modification of siRNAs may alter the efficiency of delivery methods[19]. Yet, no systematic optimization of fully chemically modified siRNA delivery to T cells has been performed to date.

Delivery routes as well as chemical modification pattern may influence the distribution of internalized siRNA between productive and non-productive pathways. For example, partially modified siRNAs delivered *via* lipid nanoparticles to liver *in vivo* are present in the cytoplasm at 50-fold excess (*i.e*. non-productive compartments) compared to siRNA guide strands loaded to Ago2 and actively driving mRNA degradation (*i.e*. productive pathway)[20]. However, when fully modified siRNAs are delivered *via* small-molecule-conjugates, 3000-fold excess of siRNA molecules can be found in cytoplasmic non-productive compartments (e.g. endosomes) compared to the Ago2-loaded siRNA guide strands [21]. The effect of the delivery method on productive siRNA uptake pathways in T cells, however, remains unexplored.

Lipid-nanoparticle-mediated delivery initially dominated the siRNA clinical development field[22] leading to the first siRNA product, Patisiran, to be approved[23]. Lipid nanoparticles enable cellular uptake of large negatively charged RNA molecules and also shield siRNAs from nucleases in the bloodstream. The latter protective function is essential when using nuclease-sensitive unmodified RNAs (such as unmodified siRNAs in initial trials or current mRNA-based vaccines) but is less so when using fully chemically modified siRNAs that are nuclease-resistant[24]. Consequently, none of the five fully chemically modified siRNAs that are currently approved rely on lipid nanoparticles for delivery[25]. In order to circumvent LNP-related toxicities, conjugate-mediated delivery has become the most widely used approach for fully chemically modified siRNA delivery. Here a small molecule, such as a sugar or a lipid is covalently attached to the siRNA passenger strand and mediates association to the lipid bilayer or a membrane-bound receptor, followed by endocytosis[26]. Even though extensively characterized in liver, LNP-mediated and conjugate-mediated siRNA delivery to T cells have not yet been comparatively analyzed.

In this study we compare lipid-nanoparticle-mediated and lipid-conjugate-mediated delivery to T cells *ex vivo* in the context of fully chemically modified siRNAs. We evaluate a widely used lipid-conjugate – cholesterol – that enables uptake to a high variety of cell types[26] including reports in T cells[17]. Furthermore, we test a recently described lipid-siRNA-conjugate – divalent myristic acid – that has shown superior accumulation in T-cell-containing organs in mice[27]. We further characterize the uptake kinetic of lipid-conjugated siRNAs to T cells and analyze the effect of the lipid-conjugates on productive uptake pathways.

Resting T cells are infamously difficult to transfect and all gene- or RNA-delivery methods to date are rather targeting activated T cells. Yet, *ex vivo* manipulation of T cells without the need of activation or cell expansion is essential for applications in allogeneic donor lymphocyte modifications, which are not routinely activated, expanded or otherwise manipulated *ex vivo*. Since both expansion and activation conditions substantially affect the efficacy o CAR-T and engineered TCR-T cell therapies[28], and shorter *ex vivo* culture duration has been shown to enhance the effector function and persistence of CAR-T cells[29], effective delivery of gene- and RNA-therapy tools to resting T cells is of great therapeutic interest. We are first to report productive siRNA uptake into resting T cells using lipid-conjugate-mediated delivery.

Since nearly all immune cell types have been shown to drive disease[30] or are explored as cell therapy candidates [31-35], modifying immune cells beside T cells using siRNAs is of high therapeutic interest. We evaluated lipid-conjugate-mediated siRNA delivery to all mononuclear immune cell types *ex vivo* (including B and NK cells, monocytes, macrophages and dendritic cells).

## Material and Methods

### PBMC isolation

Buffy coats of healthy donors were obtained from the Center of Clinical Transfusion Medicine Tuebingen. PBMCs were isolated from buffy coats by density gradient centrifugation. Briefly, the buffy coat bag was disinfected with 70% ethanol, buffy coats were then transferred to 50 ml conical tubes and diluted 1:1 with PBS (#D8537, Sigma). 35 ml of diluted buffy coat was then layered over 15 ml of Ficoll (#11768538, Fisher Scientific) and centrifuged at room temperature at 800 X g for 18 min without brake. The interphase containing PBMCs was transferred to 50 ml conical tubes, filled up to 50 ml with PBS and centrifuged at 450 X g for 5 min. Supernatant was aspirated and the pellet was resuspended in 40 ml PBS. This washing step was repeated two more times. Pellet was then resuspended in 40 ml PBS and centrifuged at 130 X g for 10 min (removal of platelets). Supernatant was then aspirated and pellet resuspended in RPMI-1640 medium (#392-0427, VWR) supplemented with 10% FBS (#11573397, Gibco), 1 % P/S (#P0781, Sigma), 25 mM Hepes (#9157.1, Carl Roth) and cells counted using trypan blue (#T8154, Sigma) and Neubauer hemocytometer (#631-0926, VWR).

### T cell activation

1,5 x10^6^ PBMCs were seeded in 1 ml RPMI-1640 medium (#392-0427, VWR) supplemented with 10% FBS (#11573397, Gibco), 1 % P/S (#P0781, Sigma), 25 mM Hepes (#9157.1, Carl Roth) in a 24-well cell culture plate (#10380932, Fisher Scientific). Loaded CD2/CD3/CD28 MACSiBeads (#130-091-441, Miltenyi Biotec) were added in a 1:2 bead-to-cell-ratio according to manufacturer’s instructions and incubated for 3 days at 37°C, 5% CO2. Medium was changed to ATC medium consisting of RPMI-1640 (#392-0427, VWR) with 10% FBS (#11573397, Gibco), 1% P/S (#P0781, Sigma), 25 mM Hepes (#9157.1, Carl Roth), 1 mM Sodiumpyruvat (#12539059, Gibco), 10 ng/ml IL-7 (#130-093-937, Miltenyi Biotec) and 3 ng/ml IL-15 (#130-093-955, Miltenyi Biotec).

### Isolation of T cells, B cells, dendritic cells, natural killer cells and monocytes

Human T cells, B cells, dendritic cells, natural killer cells and monocytes were isolated from PBMCs by negative selection and magnetic sorting using cocktails of biotin-conjugated antibodies against non-target cells (#130-096-535,Miltenyi Biotec for T cells, (#130-101-638, Miltenyi Biotec for B cells, #130-100-777, Miltenyi Biotec for dendritic cells, #130-092-657, Miltenyi Biotec for natural killer cells, (#130-096-537, Miltenyi Biotec for monocytes). A QuadroMACS™ (Miltenyi Biotec) and LS columns (Miltenyi Biotec, Bergisch Gladbach, Germany) were used according to manufacturer’s instructions. RPMI-1640 (#392-0427, VWR) with 10% FBS (#11573397, Gibco), 1% P/S (#P0781, Sigma), 25 mM Hepes (#9157.1, Carl Roth) and 1 mM Sodiumpyruvat (#12539059, Gibco) was supplemented with 10 ng/ml IL-7 (#130-093-937, Miltenyi Biotec) and 3 ng/ml IL-15 (#130-093-955, Miltenyi Biotec) for T cells, 10 ng/ml IL-4 (#130-093-920, Miltenyi Biotec) for B cells, 100 ng/ml GM-CSF (#130-093-864, Miltenyi Biotec) and 40 ng/ml IL-4 (#130-093-920, Miltenyi Biotec) for dendritic cells, 100 ng/ml IL-2 (#130-097-744, Miltenyi Biotec) for natural killer cells.

### Differentiation of monocyte-derived dendritic cells and monocyte-derived macrophages

Monocytes were isolated as described above and 3×10^6^ cells/ml were plated in a 6-well plate using RPMI-1640 medium (#392-0427, VWR) supplemented with 10% FBS (#11573397, Gibco), 1% P/S (#P0781, Sigma), 25 mM Hepes (#9157.1, Carl Roth), 1 mM Sodiumpyruvat (#12539059, Gibco) and 14,3 μM β-mercaptoethanol (#10367100, Fisher Scientific). On day 0, 2 and 4 cytokines were added as followed: 20 ng/ml IL-4 (#130-093-920, Miltenyi Biotec) and 100 ng/ml GM-CSF (#130-093-864, Miltenyi Biotec) for monocyte-derived dendritic cells and 20 ng/ml IL-4 (#130-093-920, Miltenyi Biotec) and 40 ng/ml M-CSF (#130-096-485, Miltenyi Biotec) for monocyte-derived macrophages. On day 6, cells were collected in ice cold PBS and quality control was conducted by means of flow cytometry before using cells for downstream experiments.

### Cell lines

Jurkat cells were cultured in RPMI-1640 medium (#392-0427, VWR) supplemented with 10% FBS (#11573397, Gibco), 1% P/S (#P0781, Sigma) and 25 mM Hepes (#9157.1, Carl Roth). Cells were passaged every 2 days to maintain a cell density between 1-3 ×10^6^ cells/ml. Cells were passaged every 3 days to maintain a cell density of 0.5 ×10^6^ cells/ml.

### Branched DNA assay

Cells were co-incubated with various concentrations of siRNA for 6 days at 37°C and 5% CO2 in the presence of 5% FBS. Cells were then lysed and mRNA quantification was performed using QuantiGene™ Singleplex Assay kit, (Invitrogen™, Thermo Fisher Scientific) according to manufacturer’s instructions. The following probesets were used: PPIB (siRNA target, SA-100 03, Invitrogen™, Thermo Fisher Scientific), HPRT, housekeeping in PBMC, ATC, NK, macrophages, dendritic cells SA-100 30, Invitrogen™, Thermo Fisher Scientific) and RAN (housekeeping in Jurkats, B cells, monocytes, NK92-MI, SA-15837, Invitrogen™, Thermo Fisher Scientific). The linear range of this assay was identified for each probeset in each cell type. Datasets were first normalized to housekeeping gene expression and afterwards to untreated control. Each measurement was performed in triplicates.

### Isolation of small extracellular vesicles

Small extracellular vesicles (sEVs) were isolated from umbilical cord matrix derived mesenchymal stem cells (#C-12971, PromoCell)) via differential ultracentrifugation. Briefly, cells were cultivated in Mesenchymal Stem cell Growth Medium 2 (#C-28009, PromoCell). Medium was changed to RPMI-1640 (#392-0427, VWR) without FBS or antibiotics 24 hours before sEV isolation. After 24 hours, this serum-free conditioned medium was collected and Mesenchymal Stem Cell Growth Medium 2 (#C-28009, PromoCell) added to cells for further cultivation. Conditioned medium was centrifuged at 300 X g, for 10 min at room temperature to remove dead cells. Supernatant was then centrifuged at 10,000 X g for 30 min at 4°C and filtered through a 0,2 μm membrane (Nalgene® bottle-top sterile filter, Z35223, Sigma). Small extracellular vesicles were pelleted at 100,000 X g for 1,5 h at 4°C by using Ti-45 rotor (Beckman Coulter) and 70 ml polycarbonate bottles (Beckman Coulter) in a Sorvall™ WX Ultra Series ultracentrifuge (Thermo Fisher). Pellets were pooled in 1 ml PBS and centrifuged at 100,000 X g for 1,5 h at 4°C in 1,5 ml tubes in a Ti-45 rotor (Beckman Coulter) using adapters (#11004, Beranek). Final pellet was resuspended in 100 μl PBS and frozen in a 0,1M sucrose (steril filtered, #S0389, Sigma) with a protease inhibitor cocktail (cOmplete™, Mini, #11836170001, Roche). siRNAs were co-incubated with sEVs (siRNA-to-EV ratio 10.000:1) for one hour at 37°C at 5% CO2. Then loaded sEVs were pelleted at 100,000 g for 90 minutes at 4°C and unloaded siRNA discarded. siRNA-loaded sEV pellet was then resuspended in cell culture medium and added to cells.

### Formulation of lipid nanoparticles

Lipid nanoparticles (LNPs) were formulated using NanoAssemblr® Spark™ instrument (Precision NanoSystems) and GenVoy-ILM™ T cell kit for mRNA (Precision NanoSystems) according to manufacturer’s instructions. 10 μg siRNA (around 800 pmol) were added to the hydrous phase during LNP production.

### Characterization of extracellular vesicles and lipid nanoparticles

Nanoparticle Tracking Analysis was performed to determine concentration and size distribution of sEVs and LNPs using a NanoSight NS300 (Malvern) instrument. Samples were infused using continuous syringe flow pump (infusion rate 800) and 30 second movies captured 3 times at 20°C. Detection threshold was set at 5. sEVs and LNPs were appropriately diluted in PBS prior to measurement to have 20-120 particles per frame and more than 1000 valid particles in total in order to ensure robust analysis. Zeta potential as an indicator for colloidal stability was measured by Zetasizer Nano (Malvern) with a disposable folded capillary (DTS1070) using Smoluchowski calculation. sEVs and LNPs were diluted 1:500 in PBS prior to measurement with following settings: Dispersant PBS (Viscosity: 1,0041 cP; RI: 1,330; Dielectric constant 79,0), 20°C, equilibration time 120 s and DTS 1070 cell.

### Flow cytometry

Flow cytometry was performed using a FACS Canto II instrument (BD Biosciences). Briefly, cells were harvested *via* centrifugation, then blocked with human IgG (#I4506 Sigma) for 10 minutes at 4°C, washed, then stained with live/dead stain and fluorophore-conjugated antibodies according to manufacturer’s instructions. Following antibodies were obtained from Biolegend: anti-CD3 (#300317, #317343), anti-CD4 (#300514), anti-CD8 (#300933), anti-CD19 (#363005). Antibodies against CD14 (#130-110-583) and HLA-DR (#130-111-943) were obtained from Miltenyi Biotec. Dapi (#422801, Biolegend) and Zombie NIR (#423105, Biolegend) were used to stain dead cells. PBS (#D8537, Sigma) supplemented with 0,5% BSA (#9400.1, Carl Roth) and 2 mM EDTA (#A4892,0100, PanReac AppliChem) was used as buffer. Data were analyzed with FlowJo 10.8.0 (Tree Star).

### Microscopy

Dendritic cells and T cells were isolated *via* negative selection from frozen PBMCs and a subset of T cells were activated with beads as described above. Cells were seeded (4×10^5^ dendritic cells per well, 3×10^5^ resting T cells per well and 2×10^5^ activated T cells per well) in a 96-well U-bottom plate (#10344311, Fisher Scientific) in 100 μl of cell type specific medium (see above), treated with 1 μM final concentration siRNA and incubated at 37°C, 5% CO2. 1h, 3h, 7h and 24h after siRNA treatment, cells were washed by centrifugation at 400 X g, 5 min at room temperature and resuspended in 150 μl PBS (#D8537, Sigma). Funnel, filter card (#5991022, Thermo Scientific) and microscope slide (#235504006, DWK Life Sciences) were assembled in the centrifuge (Cytospin 4, Thermo Scientific), filled with cell suspension and centrifuged at 250 rpm for 10 min. A round circle was drawn around the cell spot with a Pap-Pen (#Z377821, Sigma-Aldrich). Cells were then fixed with 4 % Paraformaldehyde (#15424389, Fisher Scientific) for 10 min at room temperature and rinsed 3 times with PBS (#D8537, Sigma). Fixed cells were stained with 1 ng/ml DAPi (#422801, Biolegend) in PBS for 10 min at room temperature in the dark and washed 3 times with PBS (#D8537, Sigma). PBS was completely removed and cells were left to dry for 5 min at room temperature. Dry cells were mounted with 1 drop of Fluoromount-G™ (#15586276, Invitrogen, Fisher Scientific), covered with cover slide (#4818602, Param GmbH) and dried overnight at room temperature in the dark. Fluorescent images were acquired using ApoTome 2 (Zeiss). Images were analyzed using Fiji (Version 2.11). Fluorescence was quantified using the pixel integrated density method.

### Statistical analysis

Silencing and uptake kinetic (flow cytometry) data was analyzed in Prism Version 10.0.1. (GraphPad). Silencing curves were fit using the “log(inhibitor) vs. response (three parameters)” function. Cellular uptake curves were fit using the “One-phase-association” function. Curves were compared using two-way ANOVA with Tukey multiple comparison tests.

## Results

### Lipid-conjugate interacts with membranes to define productive uptake of siRNAs to T cells

To compare the utility of three widespread strategies (conjugate-mediated [36], extracellular-vesicle-mediated [37] and lipid-nanoparticle-mediated [38]) to deliver siRNAs to target cells, we used a previously developed siRNA chemical structure [18]. This compound featured a 15 bp long duplex region and a fully phosphorothioated tail promoting association to membranes and proteins [19]. Furthermore, full chemical modification *via* an alternating combination of 2′-*O*-methyl and 2′-fluoro was used to confer enhanced stability [18] and immune inertness [39] to the siRNAs. The 5′-phosphate of the antisense strands was stabilized with vinylphosphonate in case of siRNAs targeting *PPIB*. 5′-phosphate stabilization does not lead to an improved silencing potency *in vitro*[40]. siRNAs were further conjugated at the 3′ end of the sense strand to either cholesterol [41] or divalent myristic acid (di-ma) [27] – two conjugates featuring nearly identical hydrophobicities [27, 41] but distinct *in vivo* distributions with divalent myristic acid driving increased siRNA association to immune organs compared to cholesterol [27]. A fluorophore (Cyanine-3) was conjugated to the 5′ end of the sense strand. Chemical modification structures of all siRNAs used in this study are depicted in supplementary Table 1. We then went on to test cholesterol- and divalent-myristic-acid (di-ma)-siRNAs in human activated T cells *ex vivo* (Figure 1). Cholesterol-siRNA and di-ma-siRNA induced similar silencing of a model target, *PPIB*, when delivered with cholesterol-siRNA showing a tendency for higher potency (Fig.1.C., p= 0.075).

**Figure 1.**
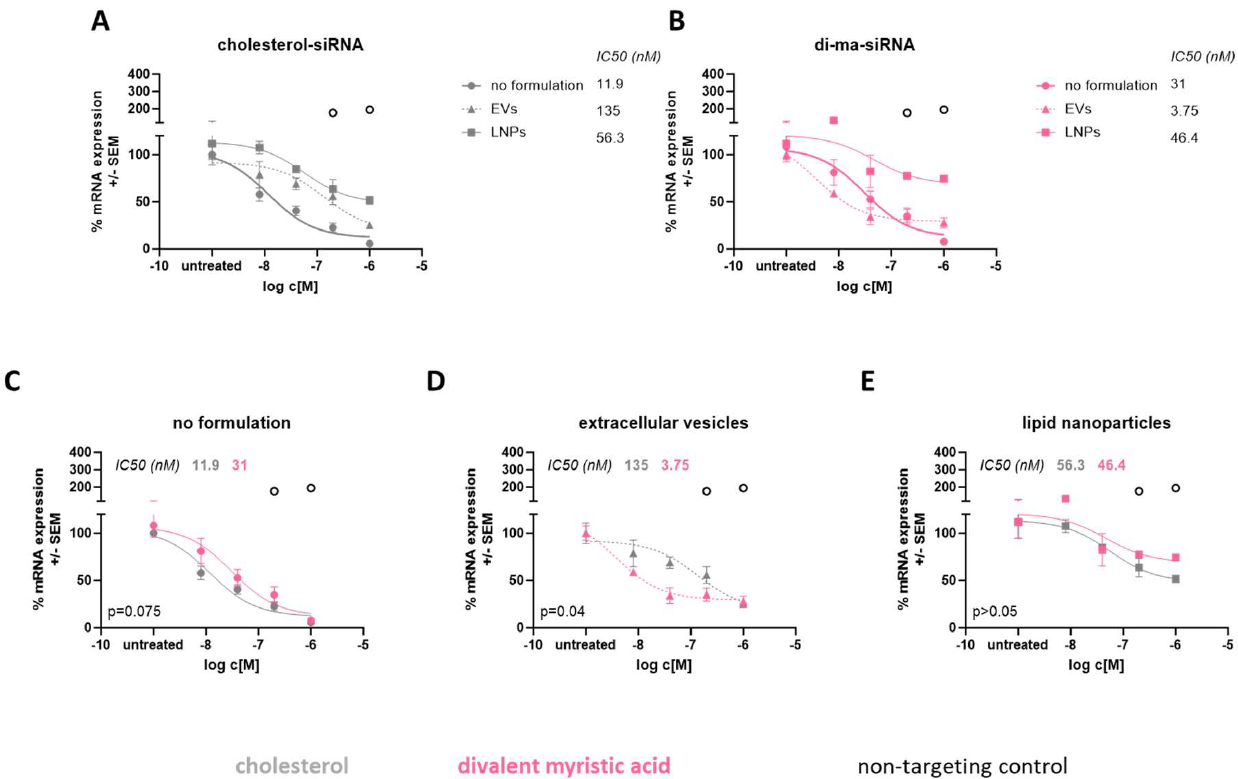
Conjugate-mediated delivery outperforms LNP- and EV-mediated delivery of siRNA to primary human T cells. T cells were isolated from PBMCs of healthy donors via negative selection and activated with CD2/CD3/CD28 beads. T cells were then treated with various concentrations of siRNA (ranging from 1 μM to 0.008 μM) conjugated to either cholesterol (A) or divalent myristic acid (B). siRNAs were either added to T cells without a formulation (C) or formulated into small EVs derived from MSCs (D) or T-cell-specific LNPs (E). Following 6 days of co-incubation, target mRNA (PPIB), and housekeeping mRNA (HPRT) expression was measured via QuantiGene Singleplex assay. N=3-6, mean±SEM

Since extracellular vesicles (EVs) have been shown to further promote siRNA delivery to hard-to-transfect cell types [42], we formulated both lipid-conjugated siRNAs to small EVs (100 nm size on average, Supp.Fig.1.A.) enriched from mesenchymal stem cell (MSC) conditioned medium. We have shown before, that hydrophobicity of lipid-conjugated siRNAs drives productive loading onto EVs [43]. Accordingly, loading efficiency of the equally hydrophobic cholesterol- and di-ma-conjugated siRNAs was indistinguishable (Supp.Fig.1.B.). Unexpectedly, EV-mediated delivery impaired the productive uptake of cholesterol-siRNA approximately 10-fold (Fig.1.A., IC50 12 nM *versus* 135 nM, p= 0.002), and failed to significantly improve silencing efficacy of di-ma-siRNA (Fig.1.B.,IC50 31 nM *versus* 3.75 nM, p>0.05). Di-ma-siRNA performed approximately 36-fold better at inducing target mRNA silencing than cholesterol-siRNA when formulated into EVs (Fig.1.D. IC50 3.7 nM *versus* 135 nM p=0.04).

With the advent of mRNA therapies, T-cell-specific LNPs are under development for RNA-delivery [44]. Here, we tested an LNP specifically developed for mRNA delivery to activated T cells *ex vivo*. We formulated cholesterol-siRNA and di-ma-siRNA into T-cell-specific LNPs (Precision Nanosystems) and after initial quality control (Supp. Fig.1. C-D.) treated activated T cells. LNP formulation impaired the productive uptake of both cholesterol-siRNA (Fig.1.A. IC50 12 nM *versus* 56nM P<0.001) and di-ma-siRNA (Fig.1.B. IC50 31 nM *versus* 46 nM p=0.0001). Interestingly, of EVs performed better than LNPs when delivering both cholesterol-siRNA (Fig.1.A. p=0.035) or di-ma-siRNA (Fig.1.B., p<0.0001).

Since lipid-conjugate-mediated delivery led to the most potent silencing, this delivery strategy was used in subsequent experiments.

### Productive siRNA uptake to T cells is defined by early uptake kinetic in T cells

In order to distinguish the role of the lipid-conjugate from the role of other hydrophobic chemical modifications (such as the single-stranded phosphorothioate tail), we compared cholesterol-siRNA and di-ma-siRNA activity with unconjugated siRNA in activated T cells (Fig.2.). The removal of the lipid conjugate led to an approximately 4.7-fold loss of activity of the siRNA (Fig.2.E., p<0.0001), and still induced a modest but significant amount of silencing (Fig.2.E., p<0.0001 unconjugated siRNA *versus* non-targeting control). These data indicate that the lipid-conjugate and other hydrophobic chemical modifications work together to mediate productive uptake of siRNAs in activated T cells. Similar results were obtained in another model of activated T cells, the IL-2 producing T cell leukemia Jurkat cell line (Fig.2.F.).

**Figure 2.**
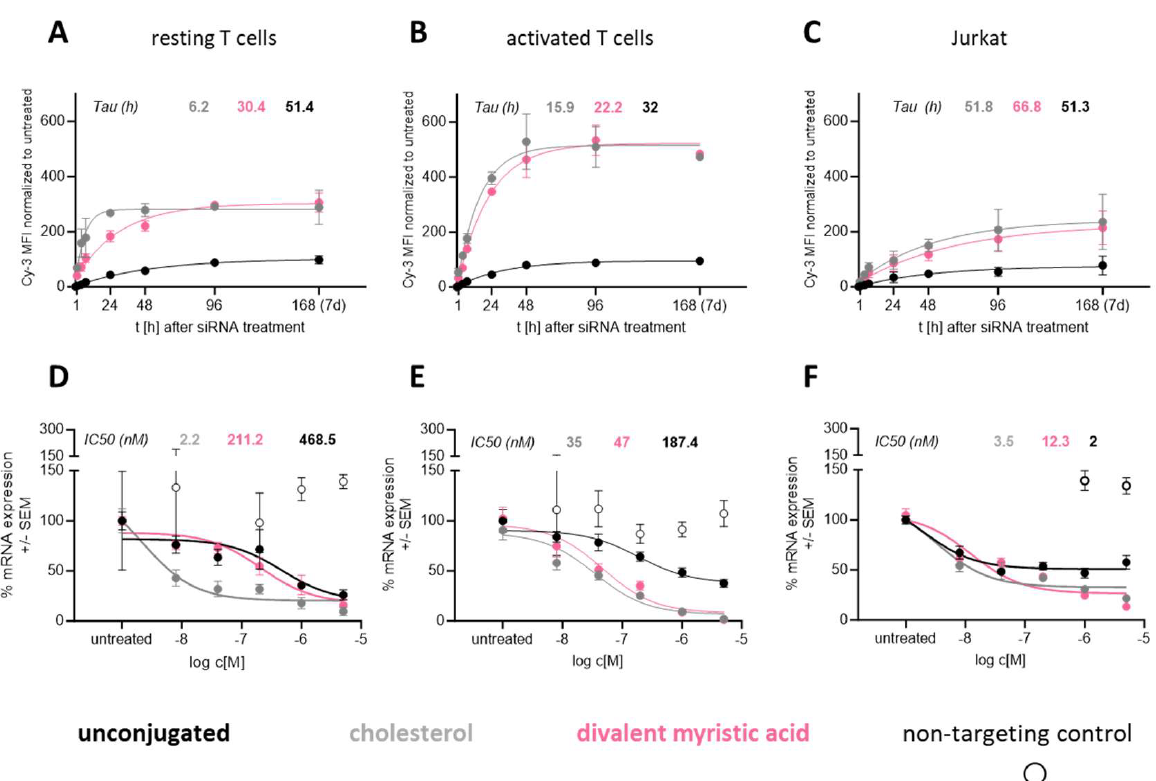
Early siRNA uptake to T cells predicts silencing efficacy. Resting (A,D), and CD2/CD3/CD28-bead-activated (B,E) T cells from healthy donors or Jurkat cells (acute T cell leukemia cell line) (C,F) were co-incubated with unconjugated (black), cholesterol-conjugated (grey) or divalent-myristic-acid-conjugated (magenta), fluorescently labeled siRNAs. For uptake experiments (A-C) cells were co-incubated with 1 μM siRNA and siRNA fluorescence (Cy3) of cells measured at 1, 4, 7, 24, 48, 96 and 168 hours post-treatment via flow cytometry. N=2, mean±SEM. Tau was calculated using the Exponential – One phase association function in Prism Version 10.0.1.In silencing experiments (D-F) cells were co-incubated with siRNAs of various concentrations (5 μM to 0.008 μM) for 6 days. Target (PPIB) and housekeeping (HPRT) mRNA expression was measured via QuantiGene Singleplex assay. Non-targeting siRNA conjugated to cholesterol (empty circles) was used as control. N=3-9, mean±SEM. IC50 was calculated via non-linear regression – (log)inhibitor – three parameters function in Prism Version 10.0.1.

Interestingly, we observed an approximately 17-fold improvement of cholesterol-siRNA silencing potency in resting T cells (Fig.2.D.) *versus* in activated T cells (Fig.2.E.). At the same time di-ma-siRNA silencing potency was 5-fold lower in resting *versus* in activated T cells (Fig.2.D-E). The presence or absence of a divalent myristic acid conjugate did not lead to a difference in productive uptake of siRNAs in resting T cells (Fig.2.D., di-ma *versus* unconjugated p>0.05). However, the presence of a cholesterol conjugate led to a 200-fold increase in silencing potency compared to other hydrophobic-modification-mediated uptake (Fig.2.D. cholesterol *versus* unconjugated p<0.0001) in resting T cells.

In accordance with the silencing data, we observed faster uptake of cholesterol-siRNA than that of di-ma-siRNA (p=0.015) or unconjugated siRNA (p<0.0001) to resting T cells (Fig.2.A.). However, this early uptake kinetics advantage disappeared after 4 days (Fig.2.A.). Yet, early (up to 2 days) uptake advantage of cholesterol-siRNA (Fig.2.A.), translated into silencing advantage (measured 6 days post-treatment) (Fig.2.D.) in resting T cells. It was somewhat surprising to see that a 3-fold higher uptake efficiency of di-ma-siRNA in comparison to unconjugated siRNA (Fig.2.A. p<0.0001) in resting T cells did not translate into an advantage in terms of silencing potency (Fig.2.D. p>0.05). Cholesterol-siRNA and di-ma-siRNA showed similar (p>0.05) uptake kinetics in activated T cells (Fig.2.B.) and in Jurkat cells (Fig.2.C.) – in accordance with silencing data (Fig.2.E and F.). Together, these data indicate (1) a membrane composition shift during T cell activation that affects interaction with hydrophobic siRNAs, and (2) the predominance of a non-productive uptake pathway for di-ma-siRNA in resting T cells.

**Figure 3.**
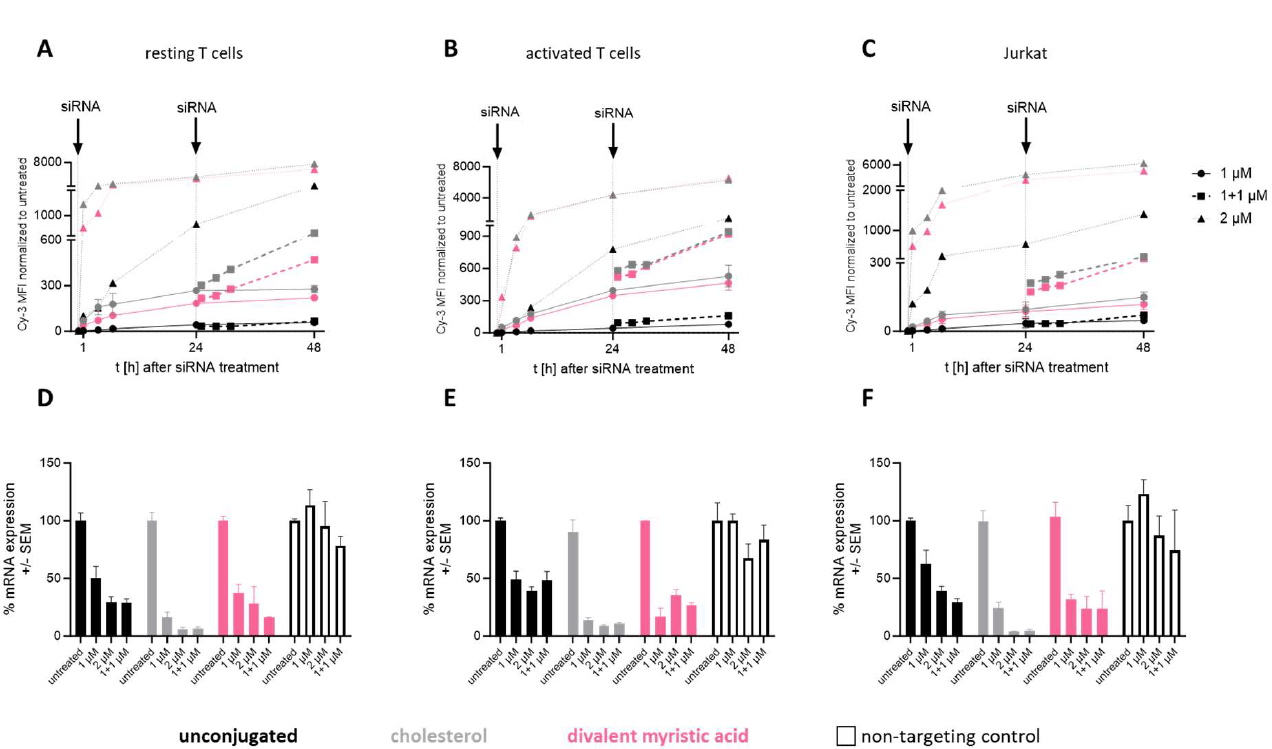
Productive uptake pathway saturates earlier than non-productive uptake. Resting (A, D), and CD2/CD3/CD28-bead-activted (B,E) primary T cells or Jurkat T cell line (C,F) were treated with either 1μM (continuous line) or 2 μM (dotted line) fluorescent siRNA conjugated to cholesterol (grey), divalent myristic acid (magenta) or unconjugated (black). In a separate set of cells, the 1 μM siRNA treatment was repeated 24 hours after the first treatment (dashed line). In uptake kinetic experiments (A-C), siRNA fluorescence associated with cells was assessed using flow cytometry at time points indicated. In silencing measurements (D-F) target (PPIB) and housekeeping (HPRT) mRNA expression was measured 6 days post-treatment via QuantiGene Singleplex assay. Non-targeting siRNA conjugated to cholesterol (empty bars) was used as control. N=3-8, mean±SEM.

Although saturation of cholesterol-siRNA uptake has been reported before to occur at around 24 hours post-treatment[19], we observed siRNA uptake saturation at a much later timepoint (4 days in activated T cells, Fig.2.B. and 7 days in Jurkat cells, Fig.2.C.). The saturation of the uptake kinetics of siRNAs could not be explained by a decrease in siRNA concentration in the medium (Supp.Fig.2.). When equimolar concentration (1μM) of siRNA was added 24 hours after the initial siRNA treatment (1 μM), a further increase in siRNA fluorescence could be observed in T cells (Fig.3. A-C, squares and dashed lines). An uptake advantage of cholesterol-siRNA *versus* di-ma-siRNA could again be observed in resting T cells (Fig.3.A.), but not in activated T cells (Fig.3.B.) or in Jurkat cells (Fig.3.C.). Interestingly, when T cells were exposed to the higher siRNA concentration (2 μM) at once, up to 200-fold increase in siRNA fluorescence could be observed associated to T cells compared to adding the same amount of siRNA in two steps, 24 hours apart (1+1μM) (Fig.3. A-C). However, above increase in uptake did not lead to an increase in silencing (Fig.3.D-F), irrespective of siRNA conjugate (cholesterol, di-ma or unconjugated) and T cell metabolic state (resting, activated or Jurkat). These data suggest that productive uptake may saturate earlier than non-productive uptake.

### Distribution of internalized siRNA depends on the lipid-conjugate and the metabolic state of T cells

We visualized intracellular localization of fluorescent siRNAs in T cells in a time course experiment using structured illumination microscopy (Fig.4.). First, we observed higher intracellular accumulation of siRNA fluorescence in activated T cells than in resting T cells (Fig.4.B *versus* A). These data were in accordance with cellular uptake data obtained *via* flow cytometry (Fig.2.). As expected, we started to see intracellular fluorescence accumulation already one-hour post-treatment consistent with fluorescence association to cells assessed *via* flow cytometry (Fig.2.B-C.).

**Figure 4.**
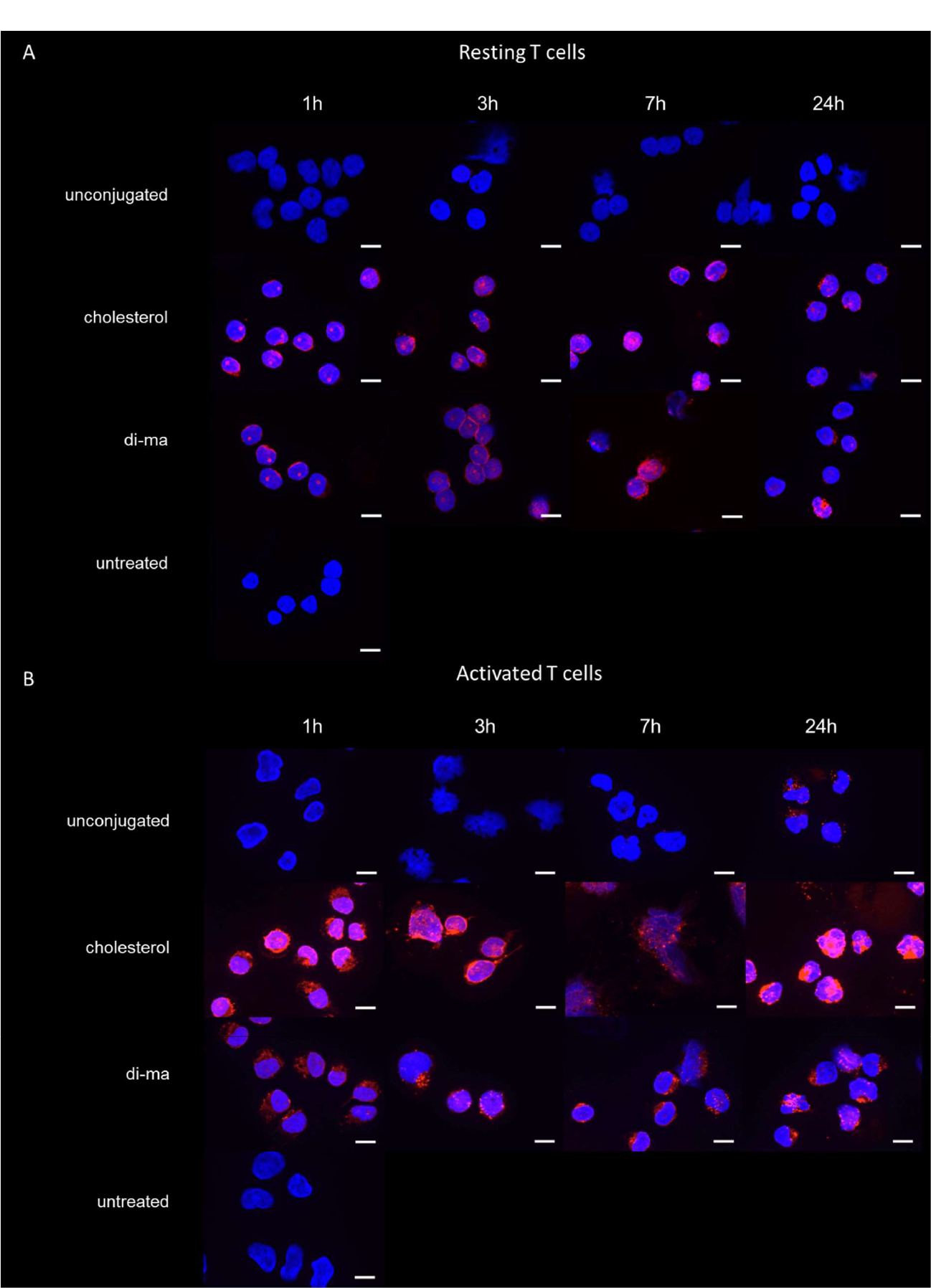
siRNA internalization depends on lipid-conjugate and metabolic state of T cells. Resting (A) and activated (B) T cells were treated with 1 μM fluorescent siRNA (magenta) for times indicated. T cells were then deposited onto a microscope slide using a cytocentrifuge and fixed with 4% PFA. Cells were stained with DAPI to visualize nuclei (blue). Images were obtained on an Apotome 2.0 (Zeiss) structured illumination microscope at 100x magnification. Bars represent 10 μm.

In resting T cells we observed perinuclear siRNA fluorescence starting at 1 hour post-treatment in both cholesterol-siRNA- and di-ma-siRNA-treated cells (Fig.4.A.). Perinuclear fluorescence showed low-grade granulation in both cholesterol-siRNA-treated and di-ma-siRNA-treated cells. We observed a modest fluorescent signal over the nucleoli starting at early timepoints, which seemed to diminish over time. In unconjugated-siRNA-treated cells, no intracellular fluorescent signal could be seen throughout tested timepoints.

In contrast, in activated T cells we observed a rather diffuse perinuclear siRNA fluorescence in cholesterol-siRNA-treated cells, while di-ma-siRNA-treated cells showed clear perinuclear fluorescent granules (Fig.4.B.). At 3 hours post-treatment intracellular fluorescent aggregates started to form in di-ma-siRNA-treated cells. Cholesterol-siRNA-treated cells showed fluorescent signal over the nuclei, especially nucleoli, and this signal seemed throughout all timepoints tested. In contrast, fluorescent signal in di-ma-siRNA-treated cells stayed in the cytoplasmic space (Fig.4.B.). Unconjugated-siRNA-treated cells showed a mild, but clearly granulated perinuclear fluorescent signal starting at 24 hours post-treatment.

Taken together, different morphology of intracellular siRNA fluorescence in resting and activated T cells as well as in di-ma-siRNA-treated and in cholesterol-siRNA-treated cells further supports the notion that (1) siRNA uptake pathways undergo a shift during T cell activation, and (2) cholesterol-siRNA and di-ma-siRNA distribute differently potentially due to different uptake mechanisms in T cells.

### Hydrophobic-interaction-mediated uptake enables functional siRNA delivery to primary mononuclear immune cells *ex vivo*

In order to characterize siRNA delivery to therapeutically relevant immune cells beside T cells, we treated a panel of mononuclear immune cells obtained from human peripheral blood with fluorescent, lipid-conjugated siRNA. We observed increasing siRNA uptake efficiency with increasing siRNA hydrophobicity in all cell types tested (Fig.5. B, D,F,I,K and M). However, silencing potency of the more hydrophobic cholesterol-siRNA and di-ma-siRNA was similar (p>0.05) than that of the less hydrophobic unconjugated siRNA in B cells (Fig.5.A.), NK cells (Fig.5.C.), monocytes (Fig.5.G.) and in macrophages (Fig.5.L.). These data suggest that productive siRNA uptake to these cell types is predominantly regulated *via* the hydrophobic phosphorothioated tail and possibly other hydrophobic chemical modifications rather than *via* the lipid conjugate. The presence of a lipid conjugate could only augment siRNA uptake *via* non-productive pathway.

**Figure 5.**
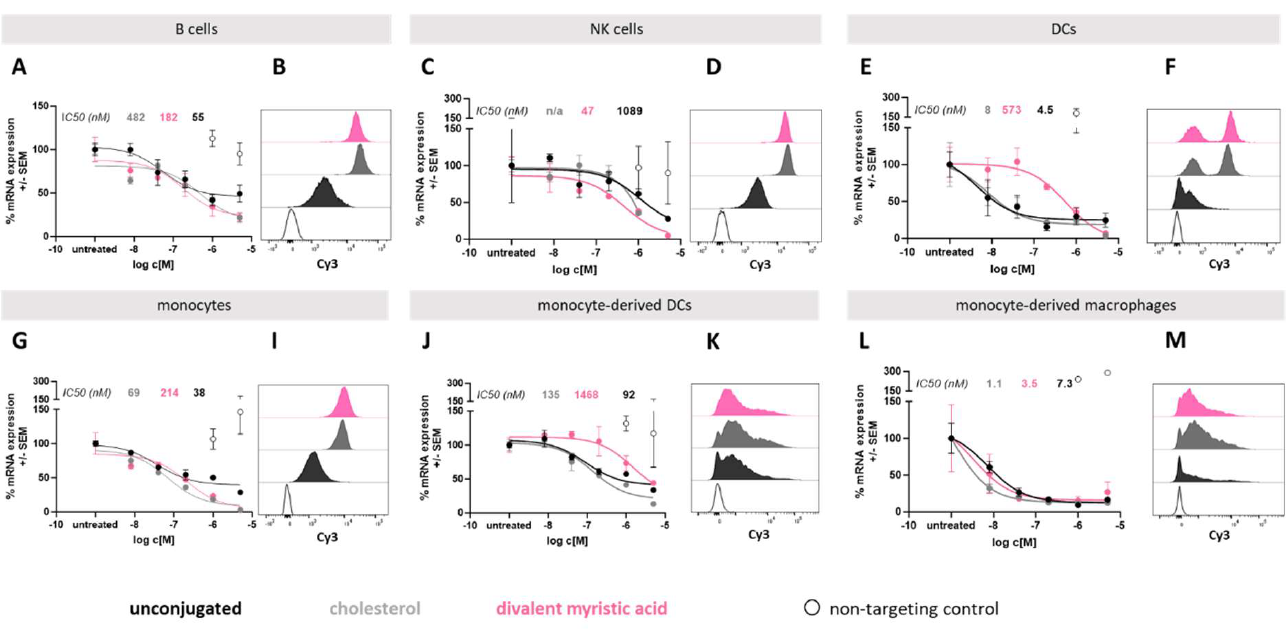
Hydrophobic interactions efficiently mediate productive uptake of siRNA to primary mononuclear immune cells. B (A-B), NK (C-D) and dendritic (E-F) cells as well as monocytes (G-I) were isolated from PBMCs of healthy donors via negative selection. Sets of monocytes were further differentiated into dendritic cells (J-K) using IL-4 and GM-CSF or macrophages (L-M) using IL-4 and M-CSF. In uptake experiments (B,D,F,I,K,M) cells were treated with 1 μM fluorescent siRNA conjugated to either cholesterol (grey) or divalent myristic acid (magenta) or unconjugated and siRNA fluorescence associated to cells measured 24 hours post-treatment via flow cytometry. In silencing experiments (A,C,E,G,J,L) cells were co-incubated with various concentration of siRNA (ranging from 5 μM to 0.008 μM) and target (PPIB) and housekeeping (HPRT or RAN) mRNA expression measured 6 days post-treatment via QuantiGene Singleplex assay.

Unexpectedly, the presence of a di-ma-conjugate on the siRNA led to a 70-fold impairment of silencing potency compared to cholesterol-siRNA (p=0.009) and a 100-fold impairment compared to unconjugated siRNA (p=0.02) in dendritic cells (Fig.5.E.). Dendritic cells in blood have two populations representing plasmacytoid and myeloid cells [45]. These two populations showed differential uptake of all siRNAs tested (Fig.5.F.). Monocytes can also be differentiated into dendritic cells, which then are more similar to myeloid than to plasmacytoid dendritic cells [45]. The presence of di-ma on siRNA also impaired silencing potency in monocyte-derived dendritic cells (Fig5.J.). Yet, in both dendritic cell models cellular uptake of di-ma-siRNA and cholesterol-siRNA was indistinguishable when assessed *via* flow cytometry (Fig.5.F. and K.). These data suggest that di-ma-conjugate actively drives siRNAs towards a non-productive uptake pathway in dendritic cells, but not in B and NK cells or monocytes. Consequently, non-productive uptake pathways must involve different signaling lipids/proteins/receptors in dendritic cells than in B and NK cells or monocytes.

All siRNAs showed very high silencing potency (IC50 < 10 nM) in macrophages (Fig.5.L.), presumably reflecting the general phagocytotic nature of this cell type.

### Phagocytosis by dendritic cells is affected *via* the presence and identity of lipid-conjugate on siRNA

To visualize productive and non-productive uptake pathways we used structured illumination microscopy to follow intracellular distribution of cholesterol-siRNA an di-ma-siRNA in dendritic cells (Fig.6.). In unconjugated-siRNA-treated cells (Fig.6.), we observed much earlier accumulation of intracellular siRNA fluorescence than in resting (Fig.4.A.) or in activated (Fig.4.B.) T cells. Rapid internalization of unconjugated siRNAs in dendritic cells may represent the primary phagocytotic function of this cell type. Similar to activated T cells (Fig.4.B.), siRNA fluorescence initially showed perinuclear granulation with increasing confluence over time in dendritic cells as well (Fig.6.). However, confluence of siRNA fluorescence seemed to be higher in cholesterol-siRNA treated cells than in di-ma-siRNA-treated cells (Fig.6.). In accordance with data obtained in activated T cells, in cholesterol-siRNA-treated dendritic cells siRNA fluorescence could also be observed over the nuclei, whereas siRNA fluorescence stayed mostly in the cytoplasm in di-ma-siRNA-treated dendritic cells (Fig.6.).

**Figure 6.**
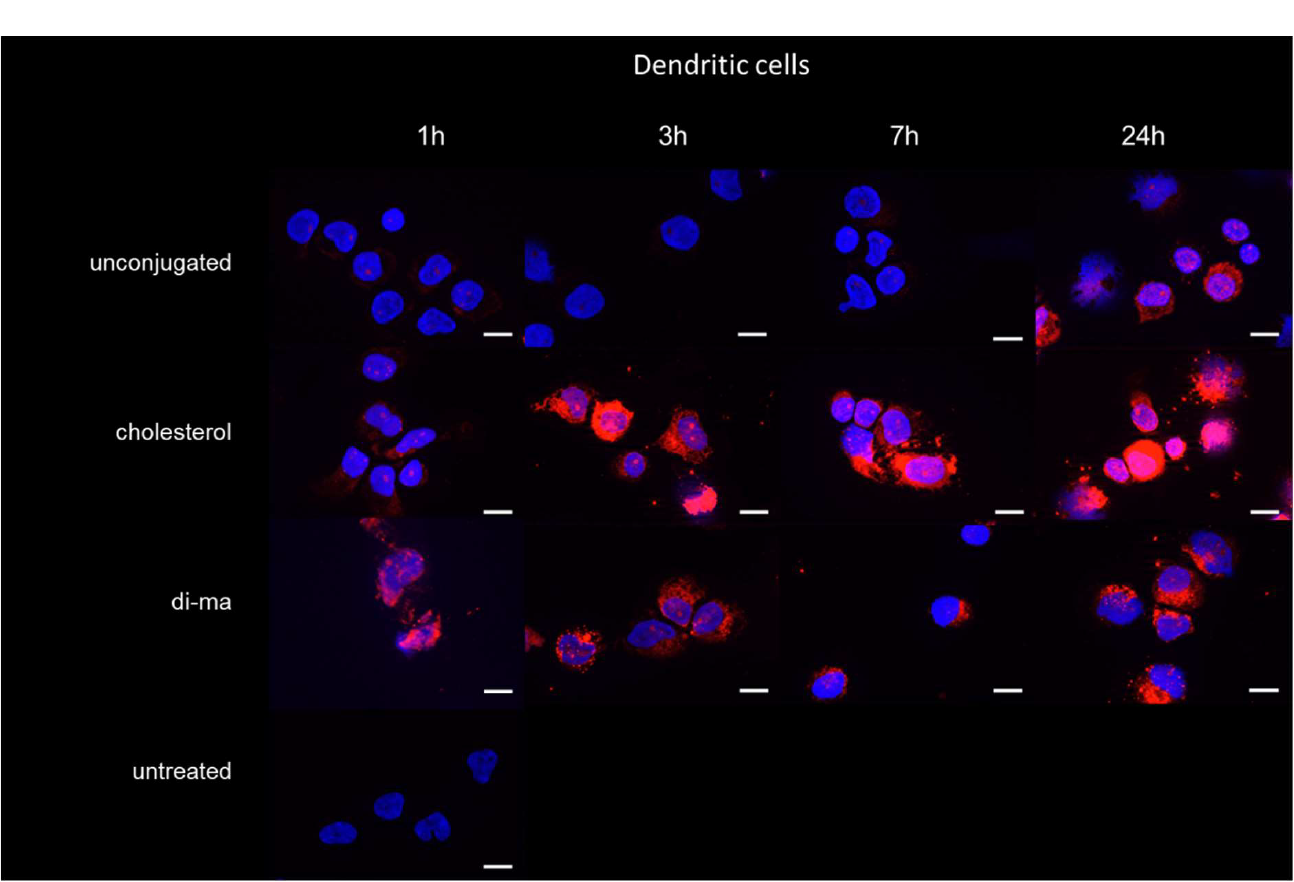
siRNA internalization in dendritic cells. Dendritic cells were isolated from PBMCs via negative selection and treated with 1 μM fluorescent siRNA (magenta) for times indicated. Dendritic cells were then deposited onto a microscope slide using a cytocentrifuge and fixed with 4% PFA. Cells were stained with DAPI to visualize nuclei (blue). Images were obtained on an Apotome 2.0 (Zeiss) structured illumination microscope at 100x magnification. Bars represent 10 μm.

Unconjugated-siRNA-treated dendritic cells showed a much lower intracellular siRNA fluorescence accumulation than cholesterol-siRNA-treated dendritic cells (Fig.6.) in accordance with cellular uptake data obtained *via* flow cytometry (Fig.5.F.) but despite leading to similarly potent silencing (Fig.5.E. and J.). Treatment with unconjugated siRNA resulted in a clear perinuclear fluorescent granulation in dendritic cells with modest confluence over time (Fig.6.). Hence, most fluorescent signal of cholesterol-siRNA in dendritic cells presumably represents non-productive uptake pathways.

## Discussion

*Ex vivo* modulation is thought to be necessary to unfold the full potential of cell-based immunotherapies [46-49]. siRNAs have been widely explored as immune cell modulators before [7-15, 17]. Here, we are first to report on side-by-side comparison of the widespread siRNA delivery strategies – conjugate-mediated, EV-mediated and LNP-mediated – to T cells *ex vivo*. In contrast to both *in vitro* and *in vivo* clinical data in hepatocytes[22, 25, 50] and in neurons[42, 51], neither EVs nor LNPs were able to enhance functional delivery of siRNAs to T cells.

LNPs used in this study have originally been developed for mRNA delivery[52]. Lipid-conjugated, fully chemically modified siRNAs are much more hydrophobic than mRNAs and therefore may enter into hydrophobic interactions with LNP components modifying LNP structure. Furthermore, productive intracellular target sites of siRNAs (RISC loading sites rough endoplasmic reticulum[53] and multivesicular body[54]) and that of mRNAs (cytoplasm) differ, therefore ideal LNP composition may also be different.

EVs are known to deliver siRNAs to cell types resistant to transfection – such as neurons – and induce enhanced silencing effects compared to both lipid-nanoparticle-mediated-delivery and lipid-conjugate-mediated delivery[24, 37, 55]. Here, we confirmed superiority of EVs over LNPs for siRNA delivery in T cells. EVs, however, failed to improve silencing activity of lipid-conjugated siRNAs in T cells, unlike in neurons[24, 37, 55]. These data suggest cell-type specific tropism of EVs. Indeed, the cell of origin[56] as well as the membrane composition[55] has been shown to affect EV homing. Even though mesenchymal-stem-cell-derived vesicles have been shown to be taken up by T cells and impede their proliferation[57] or affect their differentiation[58], lymphoid-derived vesicles may show a more effective uptake[59]. We suggest that EVs should only be used for siRNA delivery to T cells when a vesicle-intrinsic therapeutic add-on effect is expected, such as in graft-versus-host-disease[58]. Interestingly, EVs were more advantageous when delivering di-ma-siRNA than cholesterol-siRNA, even though the two compounds have nearly identical hydrophobicities[27, 60]. These data suggest lipid-specific interactions between the EV membrane and the lipid-conjugate, which may affect intracellular release and RISC loading of siRNA upon internalization.

Here, we are first to report functional conjugate-mediated siRNA uptake to resting T cells. We show differences between the early uptake kinetic and silencing potency of the equally hydrophobic compounds cholesterol-siRNA and di-ma-siRNA in resting and activated T cells. These data indicate that (1) siRNA uptake pathways are altered during T cell maturation and (2) lipid-conjugate-mediated siRNA cellular internalization not only depends on the siRNA hydrophobicity, but also on more specific interactions between the lipid-conjugate and the plasma/endosomal/lysosomal membranes and/or cytoplasmic proteins. Cellular uptake mechanisms of asymmetric siRNAs used in this study include (1) a phosphorothioate-driven mechanism similar to that of antisense oligonucleotides[19], (2) association to LDL *via* cholesterol-conjugate[61] in medium and uptake *via* LDL receptors[62], (3) hydrophobic interaction with the cell membrane[63], phagocytosis, or a combination of these. To date the uptake mechanism of di-ma-siRNA remains unknown. Since myristic acid acts as a lipid anchor to various proteins[64], it can be postulated that hydrophobic interactions contribute to cellular uptake of di-ma-siRNA. Furthermore, myristylated proteins can be bound by either heme oxygenase 2[65] or Uncoordinated-119[66], proteins playing roles in T cell signaling[65, 66] and TLR4 signaling[65]. Upregulation of these myristic acid binding proteins upon T cell activation may contribute to altered intracellular trafficking and silencing efficiency of di-ma-siRNA in activated *versus* resting T cells. Similarly, numerous cholesterol sensing or metabolizing proteins are altered during T cell activation[67], which may affect the trafficking of cholesterol-siRNA.

Even though siRNA uptake could be increased *via* repeated treatments and higher siRNA concentration, we established that productive siRNA uptake presumably saturates within the first 24 hours. Knowledge on siRNA productive uptake kinetics is essential when planning on integrating siRNA-mediated modulations into cell therapy manufacturing procedures.

Here we are first to show a comprehensive comparison of productive uptake of lipid-conjugated siRNAs to human mononuclear immune cells *ex vivo*. Data presented here demonstrate a discrepancy between cellular uptake and silencing efficiency in several immune cell types. We postulate that lipid-conjugated-siRNAs interact with the cell membrane as well as membranes of the endosomal system. This membrane-siRNA interaction will guide possible intracellular trafficking pathways towards either endosomal release and RISC loading (productive trafficking), endosomal depot formation enabling a long duration of effect[68] or degradation/inactivation (non-productive pathway). Likely, the proportion of lipid-conjugated siRNA distributed to each of these three pathways is defined *via* the siRNA-membrane interactions. Since membrane composition may largely differ between cell types [69], both the cell type and the lipid-conjugate will alter productive siRNA uptake.

Data presented here may be applied to create a library of delivery platforms to human immune cells as well as to deliver other types of oligonucleotides, such as CRISPR guide RNAs or antisense oligonucleotides (gapmer, splice-switching, RNA-editing etc.) to immune cells. Taken together, our results pave the way to future development of RNA-modified, RNA-potentiated or RNA-enabled cell therapies for cancer, autoimmune or alloimmune diseases.

## Acknowledgement

We thank Annabelle Biscans, Dimas Echeverria and Anastasia Khvorova (University of Massachusetts Medical School) for making oligonucleotides available to be used in this study. We thank the FACS Core Facility, University Hospital Tübingen for assistance with flow cytometric measurements. We thank Olga Oleksiuk (German Center for Neurodegenerative Diseases, Tübingen Site) for assistance with fluorescent imaging. We thank Philipp Schaible and Clemens Lochmann for initial assistance with QuantiGene assay establishment. We thank Emmanuelle Ribeiro for assistance with establishing PBMC isolation and primary T cell cultures. We thank Thomas Gasser (Hertie Institute, Tübingen) for access to NanoSight instrument and Philipp Bucher for initial assistance with measurements. We thank Andreas Kappler (Department of Geoscience, University of Tübingen) for access to Zetasizer and Lars Grimm for initial assistance with measurements. We thank Bruno M.D.C. Godinho and Loїc Roux for critical proof-reading of this manuscript.

This work was supported by the German Cancer Aid [70113948 to R.A.H.); and the Faculty of Medicine, University of Tübingen [473-0-0 to R.A.H., 2652-0-0 to R.A.H.].

## Author Contributions

Conceptualization A.K., R.A.H. Methodology A.K., T.R., R.A.H. Investigation A.K. Writing – Original Draft A.K., R.A.H. Visualization A.K., R.A.H. Supervision R.A.H. Project Administration R.A.H. Funding Acquisition R.A.H.

## References

1. Liu, D., et al., A practical guide to the monitoring and management of the complications of systemic corticosteroid therapy. Allergy Asthma Clin Immunol, 2013. 9(1): p. 30.

2. McMullen, J.R., et al., Ibrutinib increases the risk of atrial fibrillation, potentially through inhibition of cardiac PI3K-Akt signaling. Blood, 2014. 124(25): p. 3829–30.

3. Honigberg, L.A., et al., The Bruton tyrosine kinase inhibitor PCI-32765 blocks B-cell activation and is efficacious in models of autoimmune disease and B-cell malignancy. Proc Natl Acad Sci U S A, 2010. 107(29): p. 13075–80.

4. Giménez, N., et al., Targeting IRAK4 disrupts inflammatory pathways and delays tumor development in chronic lymphocytic leukemia. Leukemia, 2020. 34(1): p. 100–114.

5. Castillo, J.J., et al., Novel approaches to targeting MYD88 in Waldenström macroglobulinemia. Expert Rev Hematol, 2017. 10(8): p. 739–744.

6. Liu, C., et al., LINC00987 knockdown inhibits the progression of acute myeloid leukemia by suppressing IGF2BP2-mediated PA2G4 expression. Anticancer Drugs, 2022. 33(1): p. e207–e217.

7. Lokugamage, M.P., et al., Constrained Nanoparticles Deliver siRNA and sgRNA to T Cells In Vivo without Targeting Ligands. Adv Mater, 2019. 31(41): p. e1902251.

8. Freeley, M., et al., RNAi Screening with Self-Delivering, Synthetic siRNAs for Identification of Genes That Regulate Primary Human T Cell Migration. J Biomol Screen, 2015. 20(8): p. 943–56.

9. Herrmann, A., et al., CTLA4 aptamer delivers STAT3 siRNA to tumor-associated and malignant T cells. J Clin Invest, 2014. 124(7): p. 2977–87.

10. Berezhnoy, A., et al., Aptamer-targeted inhibition of mTOR in T cells enhances antitumor immunity. J Clin Invest, 2014. 124(1): p. 188–97.

11. Song, E., et al., Antibody mediated in vivo delivery of small interfering RNAs via cell-surface receptors. Nat Biotechnol, 2005. 23(6): p. 709–17.

12. Bäumer, N., et al., Targeted siRNA nanocarrier: a platform technology for cancer treatment. Oncogene, 2022. 41(15): p. 2210–2224.

13. Peer, D., et al., Selective gene silencing in activated leukocytes by targeting siRNAs to the integrin lymphocyte function-associated antigen-1. Proceedings of the National Academy of Sciences, 2007. 104(10): p. 4095–4100.

14. Kumar, P., et al., T Cell-Specific siRNA Delivery Suppresses HIV-1 Infection in Humanized Mice. Cell, 2008. 134(4): p. 577–586.

15. Iwamura, K., et al., siRNA-mediated silencing of PD-1 ligands enhances tumor-specific human T-cell effector functions. Gene Ther, 2012. 19(10): p. 959–66.

16. Mantei, A., et al., siRNA stabilization prolongs gene knockdown in primary T lymphocytes. Eur J Immunol, 2008. 38(9): p. 2616–25.

17. Ligtenberg, M.A., et al., Self-Delivering RNAi Targeting PD-1 Improves Tumor-Specific T Cell Functionality for Adoptive Cell Therapy of Malignant Melanoma. Mol Ther, 2018. 26(6): p. 1482–1493.

18. Hassler, M.R., et al., Comparison of partially and fully chemically-modified siRNA in conjugate-mediated delivery in vivo. Nucleic Acids Res, 2018. 46(5): p. 2185–2196.

19. Ly, S., et al., Single-Stranded Phosphorothioated Regions Enhance Cellular Uptake of Cholesterol-Conjugated siRNA but Not Silencing Efficacy. Mol Ther Nucleic Acids, 2020. 21: p. 991–1005.

20. Pei, Y., et al., Quantitative evaluation of siRNA delivery in vivo. Rna, 2010. 16(12): p. 2553–63.

21. Nair, J.K., et al., Impact of enhanced metabolic stability on pharmacokinetics and pharmacodynamics of GalNAc-siRNA conjugates. Nucleic Acids Res, 2017. 45(19): p. 10969–10977.

22. Jayaraman, M., et al., Maximizing the potency of siRNA lipid nanoparticles for hepatic gene silencing in vivo. Angew Chem Int Ed Engl, 2012. 51(34): p. 8529–33.

23. Adams, D., et al., Patisiran, an RNAi Therapeutic, for Hereditary Transthyretin Amyloidosis. New England Journal of Medicine, 2018. 379(1): p. 11–21.

24. Haraszti, R.A., et al., Optimized Cholesterol-siRNA Chemistry Improves Productive Loading onto Extracellular Vesicles. Mol Ther, 2018. 26(8): p. 1973–1982.

25. Brown, C.R., et al., Investigating the pharmacodynamic durability of GalNAc-siRNA conjugates. Nucleic Acids Res, 2020. 48(21): p. 11827–11844.

26. Osborn, M.F. and A. Khvorova, Improving siRNA Delivery In Vivo Through Lipid Conjugation. Nucleic Acid Ther, 2018. 28(3): p. 128–136.

27. Biscans, A., et al., The valency of fatty acid conjugates impacts siRNA pharmacokinetics, distribution, and efficacy in vivo. J Control Release, 2019. 302: p. 116–125.

28. Sudarsanam, H., R. Buhmann, and R. Henschler, Influence of Culture Conditions on Ex Vivo Expansion of T Lymphocytes and Their Function for Therapy: Current Insights and Open Questions. Front Bioeng Biotechnol, 2022. 10(886637): p. 886637.

29. Ghassemi, S., et al., Reducing Ex Vivo Culture Improves the Antileukemic Activity of Chimeric Antigen Receptor (CAR) T Cells. Cancer Immunol Res, 2018. 6(9): p. 1100–1109.

30. Mulder, W.J.M., et al., Therapeutic targeting of trained immunity. Nat Rev Drug Discov, 2019. 18(7): p. 553–566.

31. Ureña-Bailén, G., et al., Preclinical Evaluation of CRISPR-Edited CAR-NK-92 Cells for Off-the-Shelf Treatment of AML and B-ALL. Int J Mol Sci, 2022. 23(21).

32. Sloas, C., S. Gill, and M. Klichinsky, Engineered CAR-Macrophages as Adoptive Immunotherapies for Solid Tumors. Front Immunol, 2021. 12(783305): p. 783305.

33. Gabitova, L., et al., 318 Pre-clinical development of a CAR monocyte platform for cancer immunotherapy. Journal for ImmunoTherapy of Cancer, 2022. 10(Suppl 2): p. A334.

34. Liau, L.M., et al., Association of Autologous Tumor Lysate-Loaded Dendritic Cell Vaccination With Extension of Survival Among Patients With Newly Diagnosed and Recurrent Glioblastoma: A Phase 3 Prospective Externally Controlled Cohort Trial. JAMA Oncology, 2023. 9(1): p. 112–121.

35. Huang, M.N., et al., Antigen-loaded monocyte administration induces potent therapeutic antitumor T cell responses. J Clin Invest, 2020. 130(2): p. 774–788.

36. Akinc, A., et al., Targeted delivery of RNAi therapeutics with endogenous and exogenous ligand-based mechanisms. Mol Ther, 2010. 18(7): p. 1357–64.

37. Didiot, M.C., et al., Exosome-mediated Delivery of Hydrophobically Modified siRNA for Huntingtin mRNA Silencing. Mol Ther, 2016. 24(10): p. 1836–1847.

38. Zimmermann, T.S., et al., RNAi-mediated gene silencing in non-human primates. Nature, 2006. 441(7089): p. 111–4.

39. Sioud, M., Single-stranded small interfering RNA are more immunostimulatory than their double-stranded counterparts: a central role for 2’-hydroxyl uridines in immune responses. Eur J Immunol, 2006. 36(5): p. 1222–30.

40. Haraszti, R.A., et al., 5-Vinylphosphonate improves tissue accumulation and efficacy of conjugated siRNAs in vivo. Nucleic Acids Res, 2017. 45(13): p. 7581–7592.

41. Biscans, A., et al., Diverse lipid conjugates for functional extra-hepatic siRNA delivery in vivo. Nucleic Acids Res, 2019. 47(3): p. 1082–1096.

42. Didiot, M.-C., et al., Exosome-mediated Delivery of Hydrophobically Modified siRNA for Huntingtin mRNA Silencing. Molecular Therapy, 2016.

43. Annabelle Biscans, R.A.H., Dimas Echeverria, Rachael Miller, Marie-Cecile Didiot, Mehran Nikan, Loic Roux, Neil Aronin, Anastasia Khvorova, The hydrophobicity of lipid conjugated siRNAs predicts productive exosome loading. ACS Nano, 2017. To be submitted.

44. Billingsley, M.M., et al., Ionizable Lipid Nanoparticle-Mediated mRNA Delivery for Human CAR T Cell Engineering. Nano Letters, 2020. 20(3): p. 1578–1589.

45. Collin, M. and V. Bigley, Human dendritic cell subsets: an update. Immunology, 2018. 154(1): p. 3–20.

46. Schaible, P., et al., RNA Therapeutics for Improving CAR T-cell Safety and Efficacy. Cancer Res, 2023. 83(3): p. 354–362.

47. Morgan, M.A., et al., Use of Cell and Genome Modification Technologies to Generate Improved “Off-the-Shelf” CAR T and CAR NK Cells. Front Immunol, 2020. 11(1965): p. 1965.

48. Ohta, K., Y. Sakoda, and K. Tamada, Novel technologies for improving the safety and efficacy of CAR-T cell therapy. Int J Hematol, 2023. 117(5): p. 647–651.

49. Elahi, R., et al., Immune Cell Hacking: Challenges and Clinical Approaches to Create Smarter Generations of Chimeric Antigen Receptor T Cells. Front Immunol, 2018. 9(1717): p. 1717.

50. Semple, S.C., et al., Rational design of cationic lipids for siRNA delivery. Nat Biotechnol, 2010. 28(2): p. 172–6.

51. Alterman, J.F., et al., Hydrophobically Modified siRNAs Silence Huntingtin mRNA in Primary Neurons and Mouse Brain. Mol Ther Nucleic Acids, 2015. 4: p. e266.

52. McKinlay, C.J., et al., Enhanced mRNA delivery into lymphocytes enabled by lipid-varied libraries of charge-altering releasable transporters. Proceedings of the National Academy of Sciences, 2018. 115(26): p. E5859–E5866.

53. Stalder, L., et al., The rough endoplasmatic reticulum is a central nucleation site of siRNA-mediated RNA silencing. Embo J, 2013. 32(8): p. 1115–27.

54. Lee, Y.S., et al., Silencing by small RNAs is linked to endosomal trafficking. Nat Cell Biol, 2009. 11(9): p. 1150–6.

55. Haraszti, R.A., et al., Serum Deprivation of Mesenchymal Stem Cells Improves Exosome Activity and Alters Lipid and Protein Composition. iScience, 2019. 16: p. 230–241.

56. Hoshino, A., et al., Tumour exosome integrins determine organotropic metastasis. Nature, 2015. 527(7578): p. 329–35.

57. Lee, S., et al., Mesenchymal stem cell-derived exosomes suppress proliferation of T cells by inducing cell cycle arrest through p27kip1/Cdk2 signaling. Immunol Lett, 2020. 225: p. 16–22.

58. Zhang, B., et al., Mesenchymal stromal cell exosome-enhanced regulatory T-cell production through an antigen-presenting cell-mediated pathway. Cytotherapy, 2018. 20(5): p. 687–696.

59. Gutiérrez-Vázquez, C., et al., Transfer of extracellular vesicles during immune cell-cell interactions. Immunol Rev, 2013. 251(1): p. 125–42.

60. Biscans, A., et al., Hydrophobicity of Lipid-Conjugated siRNAs Predicts Productive Loading to Small Extracellular Vesicles. Mol Ther, 2018. 26(6): p. 1520–1528.

61. Osborn, M.F., et al., Hydrophobicity drives the systemic distribution of lipid-conjugated siRNAs via lipid transport pathways. Nucleic Acids Res, 2019. 47(3): p. 1070–1081.

62. Wolfrum, C., et al., Mechanisms and optimization of in vivo delivery of lipophilic siRNAs. Nat Biotechnol, 2007. 25(10): p. 1149–57.

63. Ly, S., et al., Visualization of self-delivering hydrophobically modified siRNA cellular internalization. Nucleic Acids Res, 2017. 45(1): p. 15–25.

64. Wang, B., et al., Protein N-myristoylation: functions and mechanisms in control of innate immunity. Cellular & Molecular Immunology, 2021. 18(4): p. 878–888.

65. Zhu, Y., et al., Heme Oxygenase 2 Binds Myristate to Regulate Retrovirus Assembly and TLR4 Signaling. Cell Host Microbe, 2017. 21(2): p. 220–230.

66. Gorska, M.M., et al., Unc119, a novel activator of Lck/Fyn, is essential for T cell activation. J Exp Med, 2004. 199(3): p. 369–79.

67. Bietz, A., et al., Cholesterol Metabolism in T Cells. Frontiers in Immunology, 2017. 8.

68. Badri, P., et al., Pharmacokinetic and Pharmacodynamic Properties of Cemdisiran, an RNAi Therapeutic Targeting Complement Component 5, in Healthy Subjects and Patients with Paroxysmal Nocturnal Hemoglobinuria. Clin Pharmacokinet, 2021. 60(3): p. 365–378.

69. Spector, A.A. and M.A. Yorek, Membrane lipid composition and cellular function. Journal of Lipid Research, 1985. 26(9): p. 1015–1035.

