## Supplementary Material for "Systematic optimization of siRNA productive uptake into resting and activated T cells *ex vivo*"

| target | | sequence |
|---|---|---|
| NTC | antisense | P(mU)#(fA)#(mA)(mU)(mC)(fG)(mU)(mA)(mU)(mU)(mU)(mG)(mU)#(fC)#(mA)#(fA)#(mU)#(mC)#(mA)#(fU) |
| | sense | (mU)#(mG)#(mA)(mC)(fA)(fA)(fA)(mU)(fA)(mC)(mG)(mA)(mU)#(mU)#(mA)-cholesterol |
| PPIB | antisense | V(mU)#(fC)#(mA)(fC)(mG)(fA)(mU)(fG)(mG)(fA)(mA)(fU)(mU)#(fU)#(mG)#(fC)#(mU)#(fG)#(mU)#(fU) |
| | sense | Cy3-(fC)#(mA)#(fA)(mA)(fU)(mU)(fC)(mC)(fA)(mU)(fC)(mG)(fU)#(mG)#(fA)-lipid conjugate |

| acronym | |
|---|---|
| V | 5´-(E)-Vinylphosphonate |
| # | Phosphorothioate linkage |
| m | 2´-O-methyl |
| f | 2´-fluoro |
| Cy3 | Cyanine-3 |

***Supplementary Table 1: Sequence and chemical modifications of oligonucleotides used in this study***

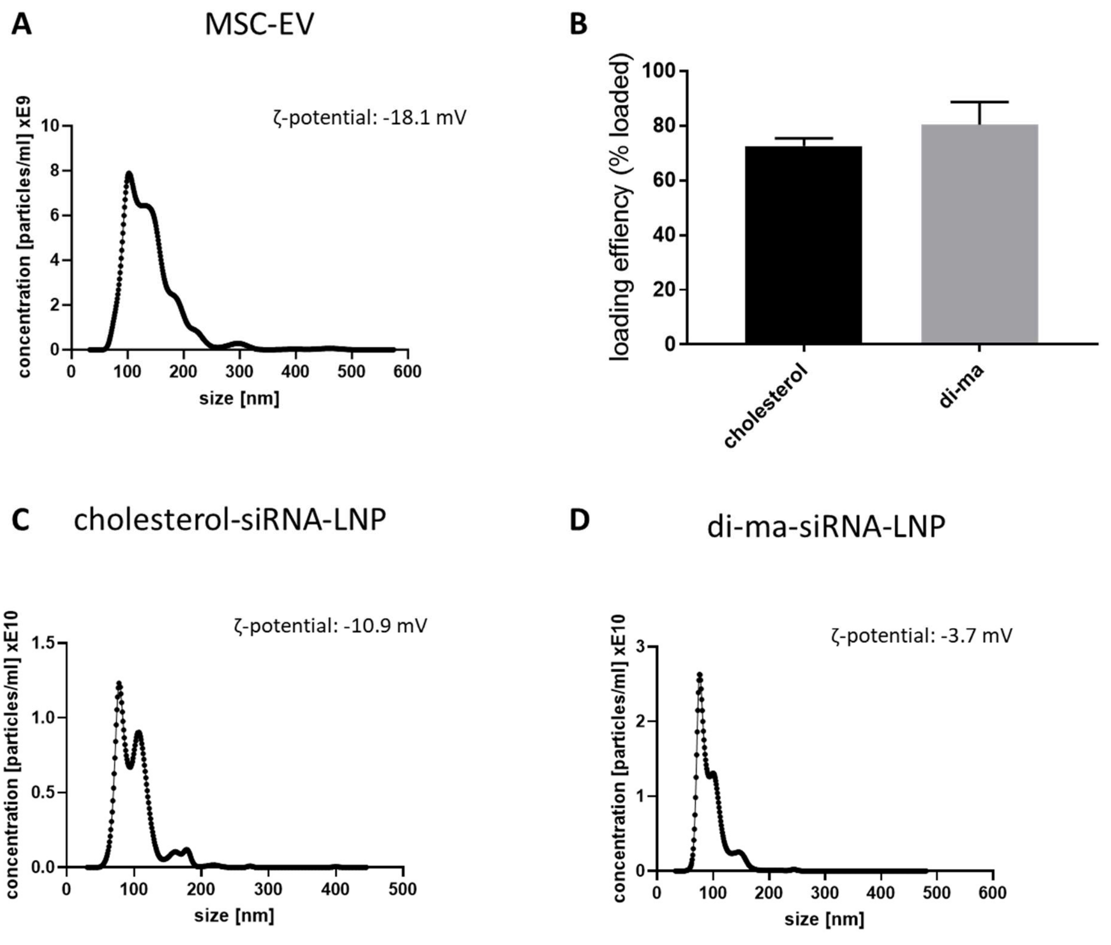


***Supplementary Figure 1 Characterization of EVs and LNPs***

*Small extracellular vesicles (EVs) were purified from conditioned serum-free media of umbilical cord, Wharton's Jelly derived mesenchymal stem cells (MSC) via differential ultracentrifugation. Size and concentration of EVs was assessed via Nanoparticle Tracking Analysis (A). Fluorescently labeled cholesterol-siRNA or di-ma-siRNA was then co-incubated with EVs at a ratio of 10.000:1 (siRNA:EV) for 1 hour ant 37℃ and the pelleted at 100.000g. Loading efficiency was calculated by comparing fluorescence in the supernatant to fluorescence in the pellet taken up in equal volume of PBS (B). Lipid nanoparticles were formulated using NanoAssemblr® Spark™ instrument (Precision NanoSystems) and GenVoy-ILM™ lipid mixture (Precision NanoSystems) according to manufacturer's instructions. 10 µg siRNA (around 800 pmol) were added to the hydrous phase during LNP production. LNP size and concentration was then characterized via Nanoparticle Tracking Analysis (C and D).*

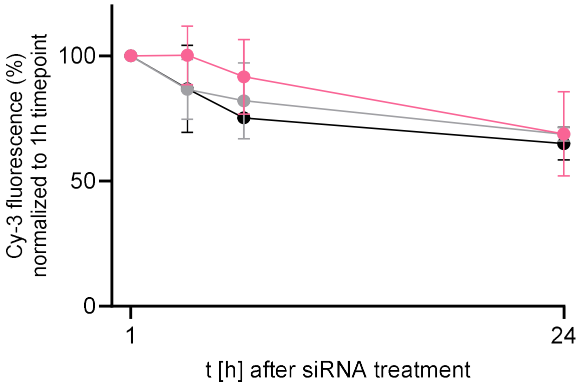


***Supplementary Figure 2 siRNA fluorescence does not substantially increase in medium upon co-incubation with cells.*** *Fluorescently labeled (Cy3) cholesterol-siRNA or di-ma-siRNA was added to Jurkat cells in round-bottom 96-well plates at a concentration of 1 µM in phenol-red-free medium (100 µl) and incubated for increasing amounts of time as indicated. Cell suspensions were then centrifuged at 500 g and samples of cell-free conditioned medium transferred to separate 96-well plates. Cy3-fluorescence was then measured in conditioned media using a fluorescent plate reader (Infinite® Pro M-Plex, Tecan). Values were normalized to the value measured at the 1-hour-timepoint. N=3, mean±SEM.*
